## Supplementary material for "Rapid Prototyping of 3D Microstructures: A Simplified Grayscale Lithography Encoding Method Using Blender": https://1drv.ms/b/s!Ap6dguNtZut4hNRyu3q8PB4E9JJRXg?e=3hXYQZ

Figures of the manuscript: "Rapid Prototyping of 3D Microstructures: A Simplified Grayscale Lithography Encoding Method Using Blender"

### Figure 1

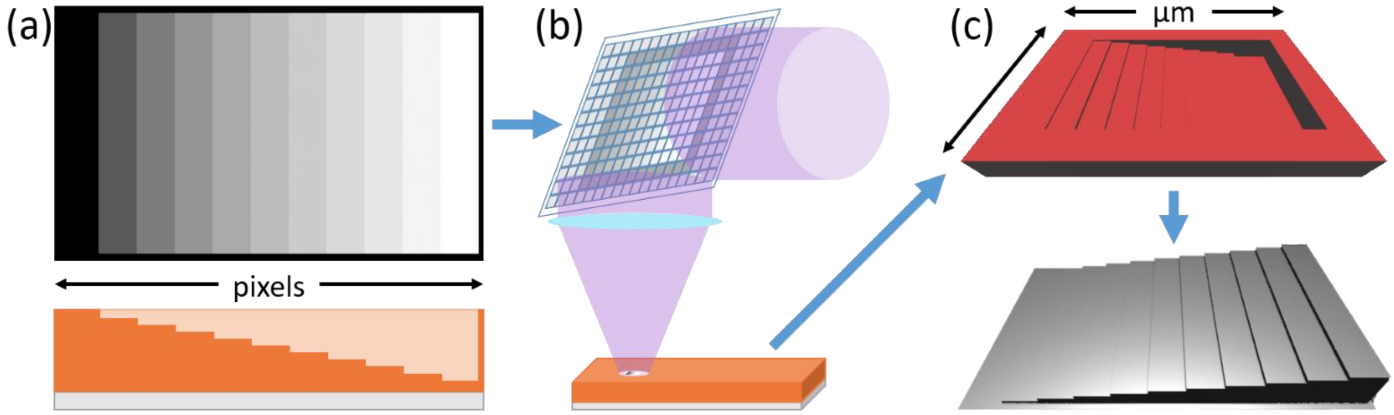


**Figure 1.** (a) Digital image in grayscale and corresponding material removal after development. Brighter gray levels correspond to higher applied power, resulting in greater material removal. (b) The image-based instructions are converted into mirror positioning on the DMD and focused onto the photoresist with appropriate intensity. (c) Representations of the expected results for the mold and soft-lithography replication.

Figure 2

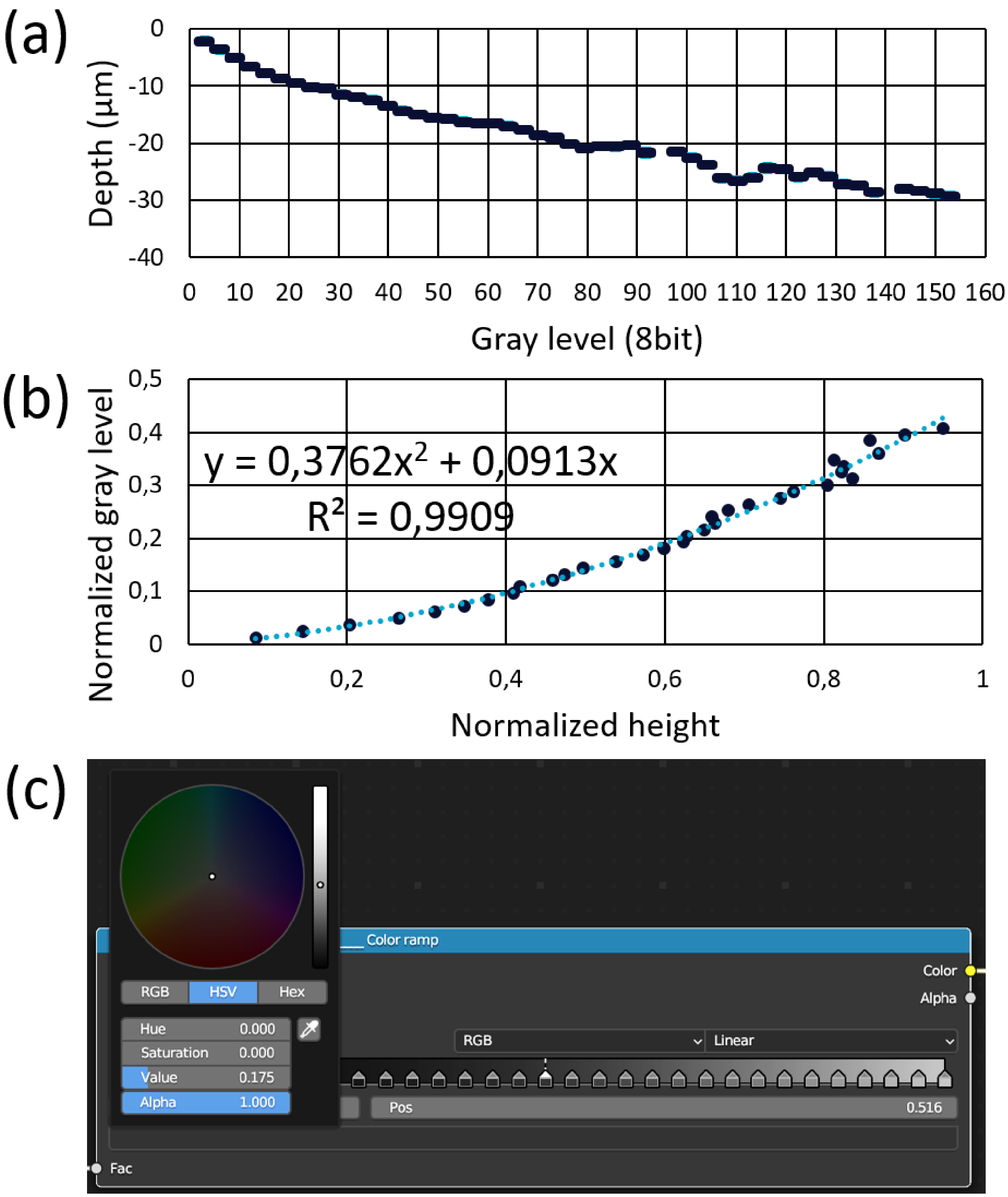


**Figure 2.** (a) Photoresist contrast curve determined by profilometry measurements following individual light power exposures corresponding to different gray levels. (b) Normalized and inverted contrast curve with fitted data. (c) Blender color ramp configured based on the calibration curve.

Figure 3

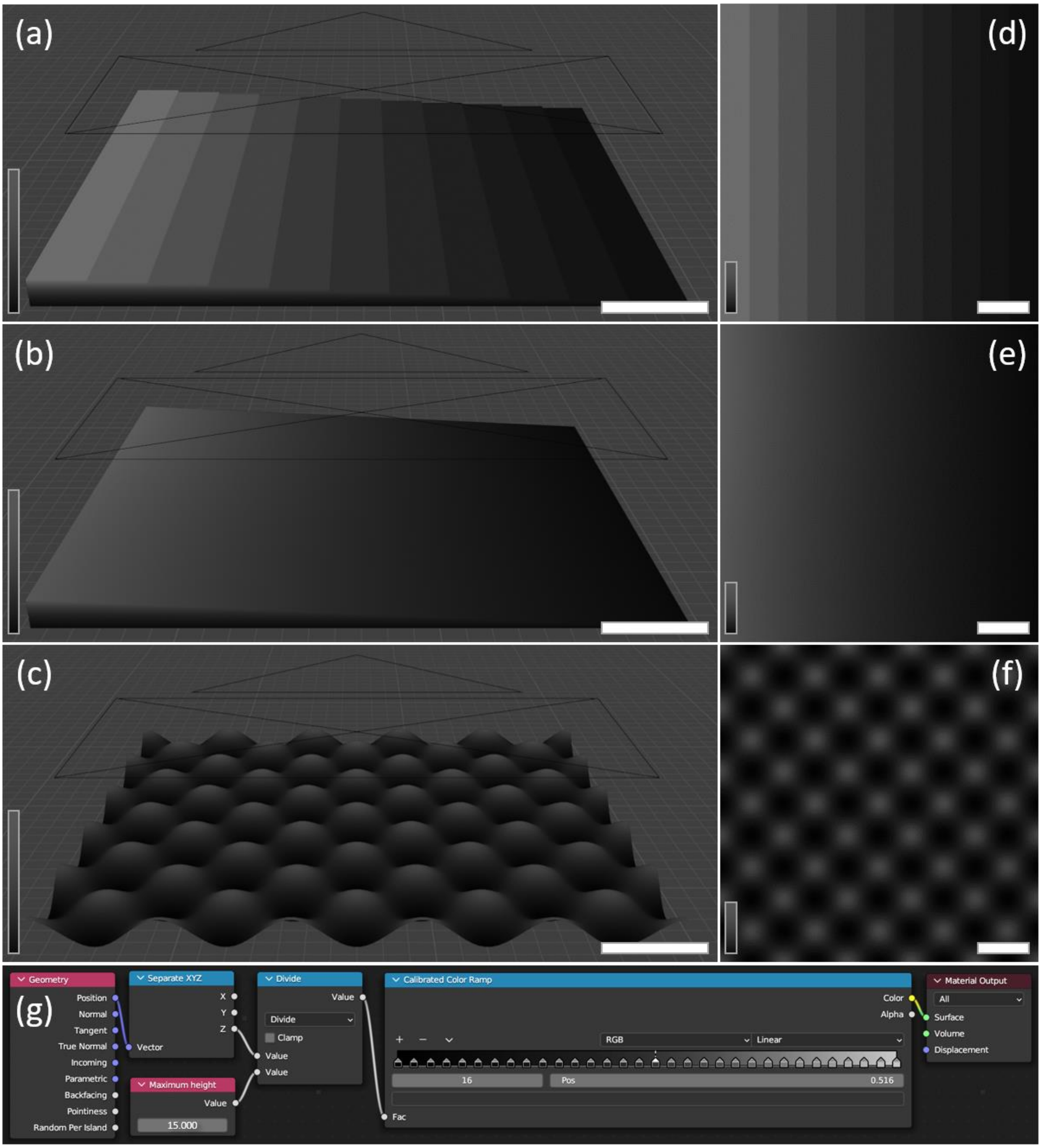


**Figure 3.** Three-dimensional designs of (a) stairs, (b) ramp, and (c) sinusoidal patterns, shaded according to depth (Z-axis). Scale bars represent 40 µm. (d, e, f) Rendered images of each model. Scale bars represent 40 µm. (g) Example of shading node setup for calibrating a maximum depth of 15 µm. Color is assigned based on Z-axis position, with black representing 0 µm and light gray representing 15 µm of material removal.

Figure 4

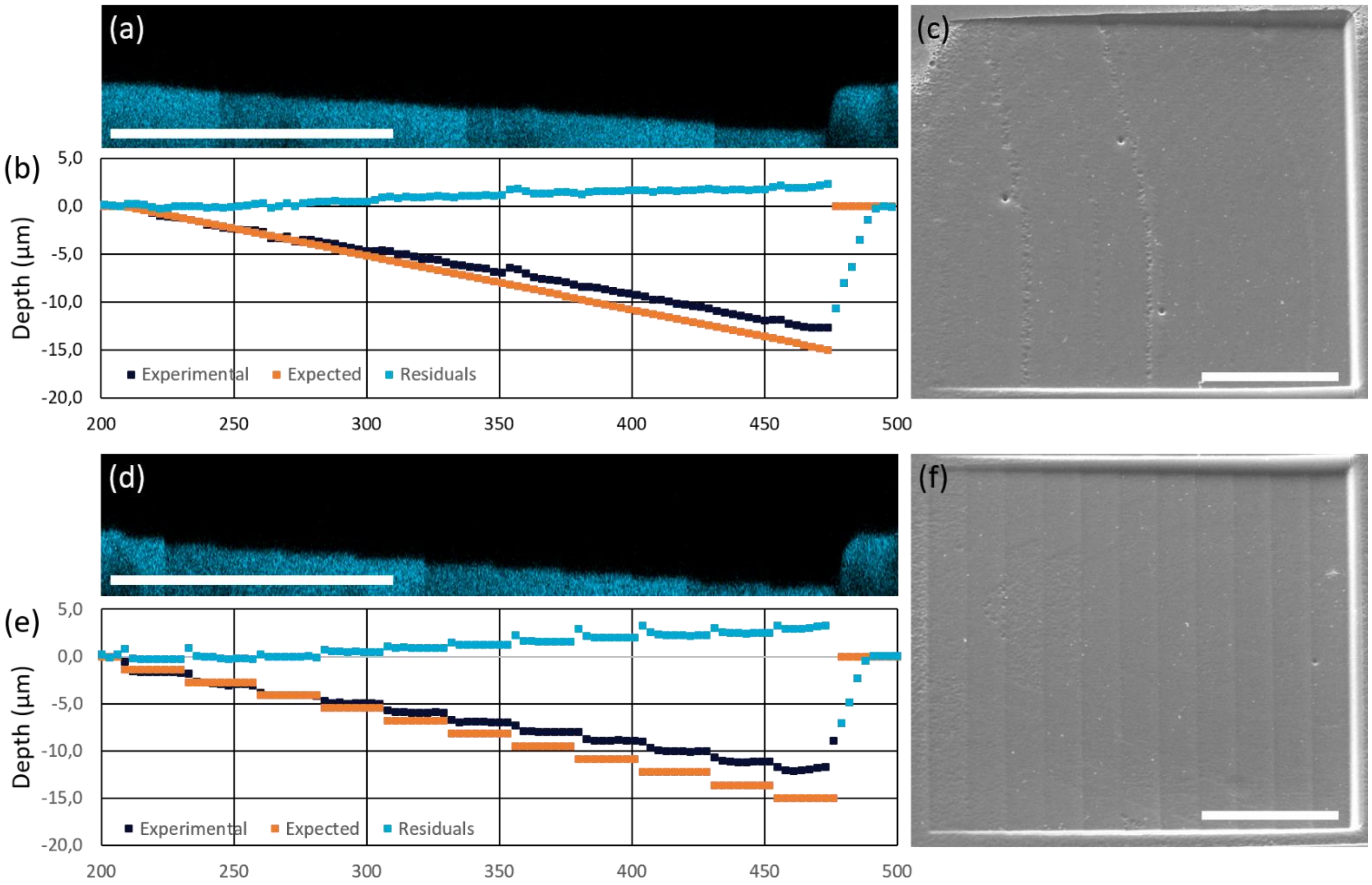


**Figure 4.** (a, d) Confocal images of exposed ramp and stair designs, respectively, based on autofluorescence. (b, e) Comparison of experimental data, expected results, and their corresponding residuals for ramp and stair designs, respectively. (c, f) SEM images of exposed ramp and stair designs tilted at 30°. Scale bars: 100 µm.

Figure 5

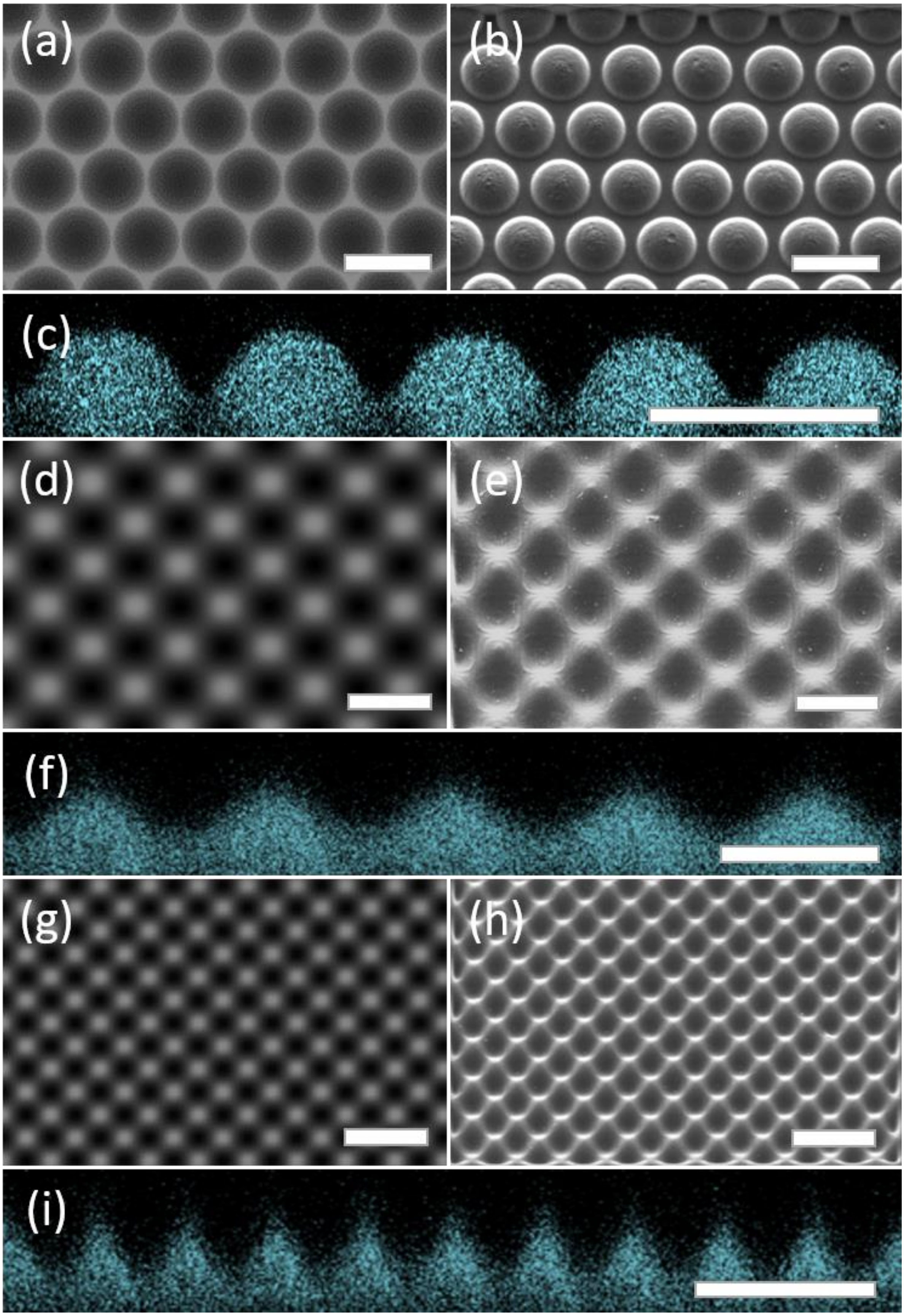


**Figure 5.** (a, d, g) Rendered designs of microlenses, sinusoidal surfaces, and doubled-frequency sinusoidal surfaces, respectively, generated using the proposed method. (b, e, h) Corresponding SEM images of developed surfaces tilted at 30°. (c, f, i) Corresponding confocal microscopy images of developed surfaces in orthogonal view. Scale bars: 50 µm.

Figure 6

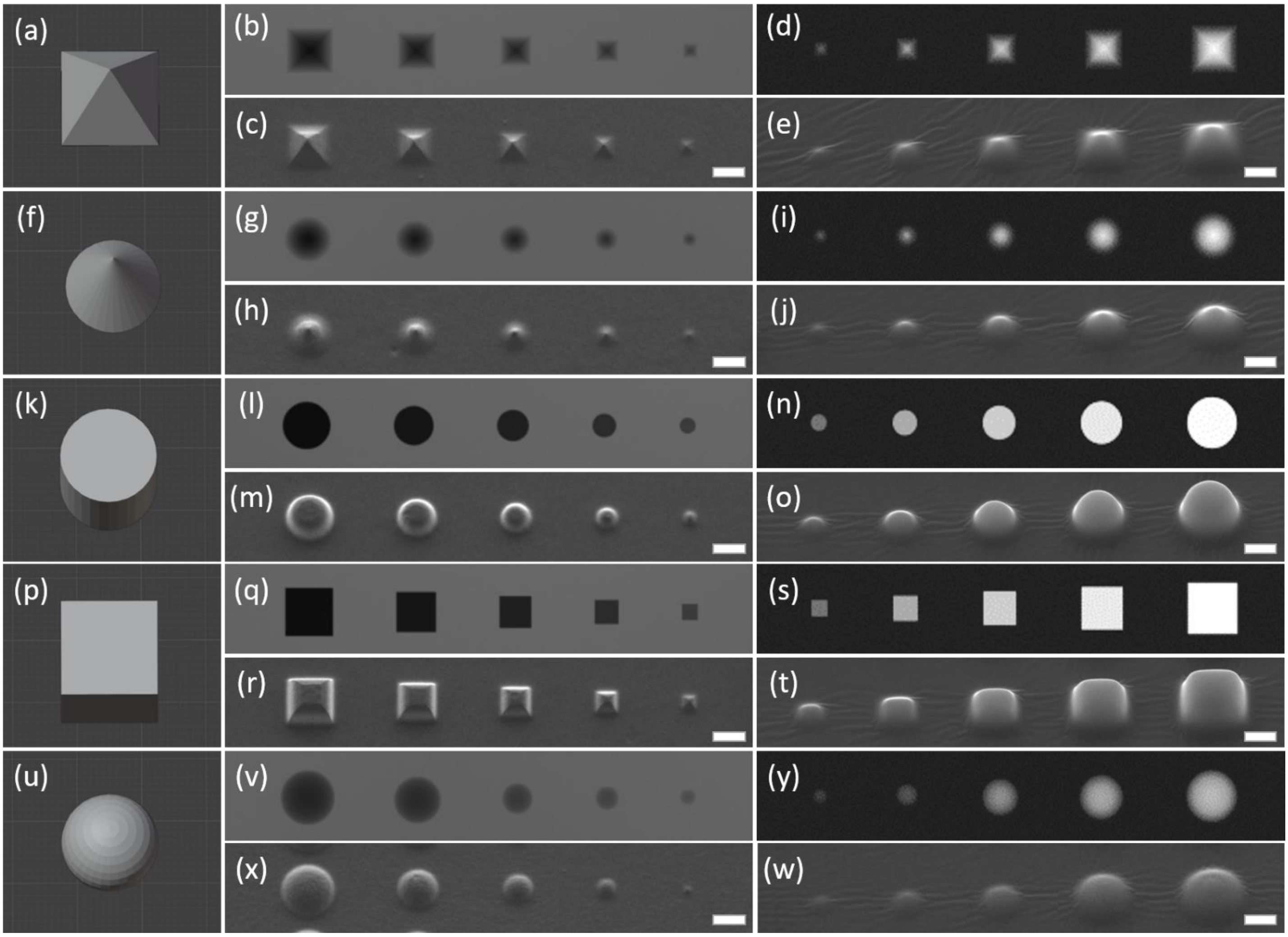


**Figure 6.** (a-e) Pyramid, (f-j) cone, (k-o) cylinder, (p-t) cube, and (u-w) hemisphere. (a, f, k, p, u) 3D designs. (b, g, l, q, v) Generated grayscale masks for direct lithography (5-15 µm). (c, h, m, r, x) SEM images of fabricated structures (20° tilt). (d, i, n, s, y) Generated grayscale masks for PDMS molding (5-15 µm). (e, j, o, t, w) SEM images of PDMS replicates (20° tilt). Scale bars: 10 µm.
